# Real-time Malaria Detection in the Amazon Rainforest via Drone-Collected eDNA and Portable qPCR

**DOI:** 10.1101/2025.06.16.660023

**Authors:** Yin Cheong Aden Ip, Luca Montemartini, Jia Jin Marc Chang, Andrea Desiderato, Nicolás D. Franco-Sierra, Christian Geckeler, Mailyn Adriana Gonzalez Herrera, Michele Gregorini, Meret Jucker, Steffen Kirchgeorg, Martina Lüthi, Frederik Bendix Thostrup, Guglielmo Murari, Marina Mura, Paola Pulido Santacruz, Florencia Sangermano, Tobias Schindler, Claus Melvad, Stefano Mintchev, Kristy Deiner

## Abstract

Emerging zoonotic parasites such as *Plasmodium* spp. in remote ecosystems demand rapid, field-deployable surveillance. We developed a drone-based environmental DNA (eDNA) sampling approach integrated with a portable quantitative PCR (qPCR) assay for in situ detection of *Plasmodium* DNA. Field trials confirmed malaria parasite DNA in canopy swabs, coinciding spatially with acoustic detection of howler monkeys (*Alouatta* sp.), suspected reservoir hosts. No *Anopheles* mosquitoes were captured in simultaneously deployed insect traps. The end-to-end workflow, from drone deployment to diagnostic readout, averaged 1.5 hours per assay, required only basic mobile-laboratory infrastructure, and bypassed cold-chain logistics. This rapid, non-invasive method delivers actionable surveillance data in real-time, enabling near-real-time trigger points for vector-control interventions and clinical response in remote settings. By merging environmental genomics with portable diagnostics, we establish a scalable One Health framework adaptable to diverse vector-borne parasites and ecosystems, with potential for early detection and spillover risk mitigation in under-monitored, high-risk environments.

## 1. Introduction

Emerging zoonotic parasites, particularly those transmitted between wildlife and humans, account for over 60% of emerging infections (Jones et al., 2008), underscoring the need for enhanced surveillance strategies to mitigate future outbreaks (Chakraborty et al., 2024; Ip et al., 2023). The increasing frequency and magnitude of zoonotic spillovers, driven by anthropogenic activities such as deforestation, wildlife trade, and climate change, further amplify the urgency for integrated surveillance strategies (Saba Villarroel et al., 2023). Recent advances in molecular tools, such as real-time PCR, metagenomics, and environmental genomics, have enhanced our ability to detect and respond effectively to emerging novel infectious agents in wildlife, offering critical insights into the dynamics of infectious diseases (Roberts et al., 2024). Expanding surveillance capabilities is imperative, particularly in biodiverse, resource-limited, and even remote regions where the ecological interactions between parasites, pathogens, hosts, and vectors remain complex and poorly understood (Le Breton et al., 2025).

Malaria is a prime example of a vector-borne disease that continues to pose a significant global health threat (Fikadu et al., 2023). Despite extensive eradication efforts, malaria remains prevalent, with an estimated 263 million cases and 597,000 deaths reported in 83 countries worldwide in 2023 (World Health Organization [WHO], 2024). The disease is caused by protozoan parasites of the genus *Plasmodium*, of which five species (*P. falciparum, P. vivax*, *P. ovale, P. knowlesi* and *P. malariae*) are responsible for causing disease in humans through transmission of bites of infected *Anopheles* mosquitoes. Timely identification of silent *Plasmodium* reservoirs is critical to inform vector-control operations and prevent recrudescence, particularly in hard-to-reach forested areas where routine entomological surveillance is logistically constrained (Le Breton et al., 2025). In Latin America, most of the malaria burden is attributable to *P. vivax* (Recht et al., 2017). Although the global burden of malaria has decreased in many regions, it continues to cause significant morbidity, mortality, and economic loss, particularly in tropical and subtropical areas such as South America (Moreno-Gutierrez et al., 2020; WHO, 2024). The situation is especially critical in developing countries, where malaria prevention and control measures still face significant challenges (Recht et al., 2017; Gunderson et al., 2020; Monroe et al., 2022; Grillet et al., 2021).

The Amazon rainforest, a global biodiversity hotspot, presents a unique opportunity to study malaria dynamics due to its complex ecosystem and the reservoir potential of diverse host species. The Amazon basin has historically been a malaria-endemic region, with the state of Amazonas, Brazil, reporting one of the highest infection rates (Lana et al., 2021). *Anopheles darlingi* is the primary vector, breeding in water bodies along river margins (Hiwat and Bretas, 2011). However, transmission dynamics are not uniform across the basin, with some areas being more prone to unstable or epidemic malaria and other regions experiencing a more stable or endemic transmission pattern (e.g. Suárez-Mutis et al., 2007). The Amazon’s distinctive hydrology—comprising blackwater, whitewater, and clearwater rivers—plays a crucial role in shaping mosquito habitats and breeding sites (Fonseca et al., 2022). Blackwater rivers, such as the Rio Negro, are known for their high acidity and low nutrient content, creating environments less favorable for mosquito breeding (Fonseca et al., 2023). Likewise, deforestation exerts a significant influence on mosquito habitat suitability and malaria transmission potential in the Brazilian Amazon, but the relationship is complex. In some contexts, higher forest cover is associated with increased malaria risk due to greater vector habitat availability and human-vector contact at forest edges (Arisco et al., 2024), while in others, deforestation facilitates mosquito proliferation by creating sunlit breeding sites and altering local ecology (Gonzalez Daza et al., 2023; Vittor et al., 2006). Consequently, some regions of the Amazon exhibit relatively low mosquito densities, which may contribute to lower malaria transmission despite the presence of malaria parasites in potential reservoir hosts.

Given the need for effective surveillance in these ecologically diverse yet remote regions, our study employs drone-assisted environmental DNA (eDNA) sampling methods combined with portable molecular tools to enhance parasite detection and monitoring capabilities (Ip et al., 2024; Aucone et al., 2023; Kirchgeorg et al., 2024; Koubínová et al., 2025). Drones enable access to hard-to-reach areas, such as the forest canopy, where traditional sampling methods are often impractical or unsafe (Cannon et al., 2021). In addition, eDNA sampling provides a non-invasive approach to detect pathogens and parasites and monitor biodiversity by analyzing genetic material shed by organisms into the environment. Portable quantitative PCR (qPCR) assays, which require minimal reliance on cold chains and sophisticated equipment, enable real-time surveillance, improving the efficiency of detection strategies (Chang et al., 2020; Jeunen et al., 2022; Roy et al., 2022). By leveraging the combination of these innovative techniques, we can conduct large-scale, remote surveillance and gather real-time comprehensive data on parasite presence and species interactions.

This study serves as proof of concept for the eDNA detection of *Plasmodium* spp. parasites, the causative agents of malaria, and their ecological relationships with potential reservoir hosts and mosquito vectors. While howler monkeys (*Alouatta seniculus*) have been implicated as key reservoirs in some regions (Abreu et al., 2019), other non-human primates, and potentially even additional wildlife, may also contribute to parasite maintenance (Hiwat & Bretas, 2011). Understanding these broader host-vector-parasite interactions is essential for anticipating zoonotic spillover risks and informing public health surveillance strategies. By combining remote eDNA sampling with drone deployment and portable parasite assays, we demonstrate a scalable, field-adaptable framework for investigating malaria dynamics in remote and biodiverse environments. This integrative approach not only enhances early detection of emerging infectious diseases but also supports One Health strategies that account for the interdependence of environmental, animal, and human health (Zinsstag et al., 2021).

## 2. Methods

### 2.1 Study area and tree canopy eDNA sample collection

The study was conducted at the XPRIZE competition test site within the Amazon rainforest in July 2024, a region characterized by its high biodiversity and complex ecological interactions. Sampling sites were located within a 1 km² grid (0.75 × 1.34 km) centered at 2.9671° S, 60.7432° W (WGS 84) on the banks of the Rio Negro (Figure 1A)—a blackwater river known for its high acidity and low nutrient levels, which creates a natural barrier to Anopheles mosquito breeding (Fonseca et al., 2023). Access to genetic resources was registered in SISGEN under XPRIZE Rainforest - Equipe ETH BiodivX - Cadastro n° A0FB047. In accordance with XPRIZE regulations, the biodiversity survey was conducted remotely using robotic platforms. Drones were employed for aerial imaging, surface eDNA collection from tree canopies, water eDNA sampling, and the deployment of canopy rafts—sensing platforms positioned above the canopy to collect overnight data. While the present report focuses on surface eDNA collection and the data acquired with the canopy rafts, a comprehensive description of the full robot-based survey system is provided in Geckeler et al. (2025).

**Figure 1.**
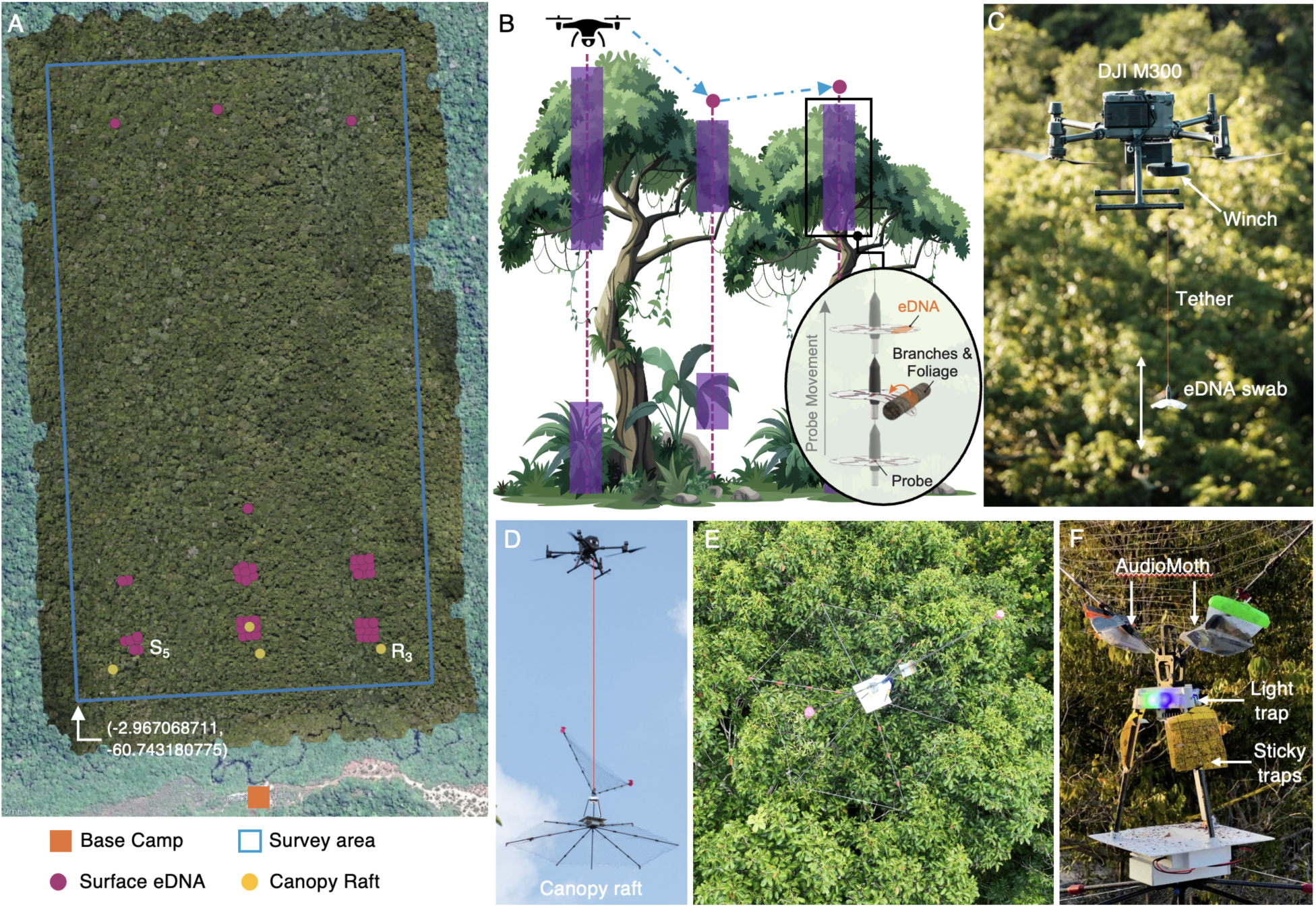
Overview of the survey technology and methodology. (A) Map of the 100-hectare survey site (blue frame) and the base camp used for drone deployments. Purple dots indicate locations where the eProbe was lowered into the canopy to collect eDNA, and yellow dots mark the deployment sites of the canopy raft. (B, C) Operating principle of the eProbe: swabs are lowered into the canopy from a hovering drone using a robotic winch system. Surface eDNA is collected upon contact with vegetation. (D) Canopy raft transported by a drone for deployment atop the tree canopy (E). (F) The canopy raft integrates sensing and sampling instruments, including an AudioMoth acoustic recorder, light trap, and three sticky traps. This image was taken when the canopy raft was brought back for the overnight sampling highlighting the insects’ specimens on the sticky traps.

Drones equipped with custom-designed eDNA swabbing mechanisms (i.e., eProbes) were deployed to collect eDNA from the forest canopy layer, following the protocol outlined by Kirchgeorg et al. (2024). Briefly, drones were flown over selected trees at the XPRIZE test site, and a swab-based eDNA collection device, attached to a robotic winch, was lowered into the canopy with a tether. The eProbe made physical contact with branches and foliage during both descent and ascent phases, facilitating the collection of eDNA, including bodily fluids, feces, and other materials shed by organisms inhabiting or interacting with the canopy. This method allowed for the capture of eDNA from various vertebrates, such as arboreal primates and birds, as well as invertebrates like insects. In total, 10 canopy swabs and two field blanks were collected during a systematic survey of a 100-hectare area of rainforest (Figure 1 A-C).

To minimize contamination risks and ensure data integrity, strict protocols were followed in the field. Before and after each use, the eProbe housing was cleaned with 10% bleach and wiped with deionized water, and a new pre-sterilized and individually packaged canopy swab was attached to the winch system. The canopy swabs were dampened with 10 mL of individually packaged molecular grade DNA-free water before deployment to facilitate particle attachment to the material. Upon return from deployment, canopy swabs were placed into sterile 50 mL Falcon tubes immediately after retrieval and eDNA extraction was performed immediately in situ, using a modified and rapid protocol optimized for eDNA extraction tailored for field conditions (Kirchgeorg et al., 2024). The resultant 5 mL volume of DNA extract was subsequently concentrated twice using an Amicon® Ultra Centrifugal Filter, 30 kDa MWCO tube (following manufacturers’ protocol) to a final 1 mL volume. The concentrated DNA was immediately subjected to further qPCR analysis.

### 2.2 Remote *Plasmodium* spp. qPCR detection

The canopy eDNA extracts were screened for *Plasmodium* spp. using a portable Diaxxo AG qPCR system (Figure 2). The multiplex qPCR assay was designed by Diaxxo AG, which simultaneously targets five human-infecting *Plasmodium* species (*P. falciparum, P. vivax, P. malariae, P. ovale, and P. knowlesi*) (Kantele et al., 2011). Of these species, only *P. knowlesi* and *P. vivax* have primate reservoirs in nature. The assay’s lower limit of detection, determined with PlasmoPod, was 0.2 parasites/µL (Bechtold et al., 2023; Stabler et al., 2025). All reactions were run in duplicates to ensure reliability, and the entire amplification protocol took under 40 minutes, facilitating rapid field deployment.

**Figure 2.**
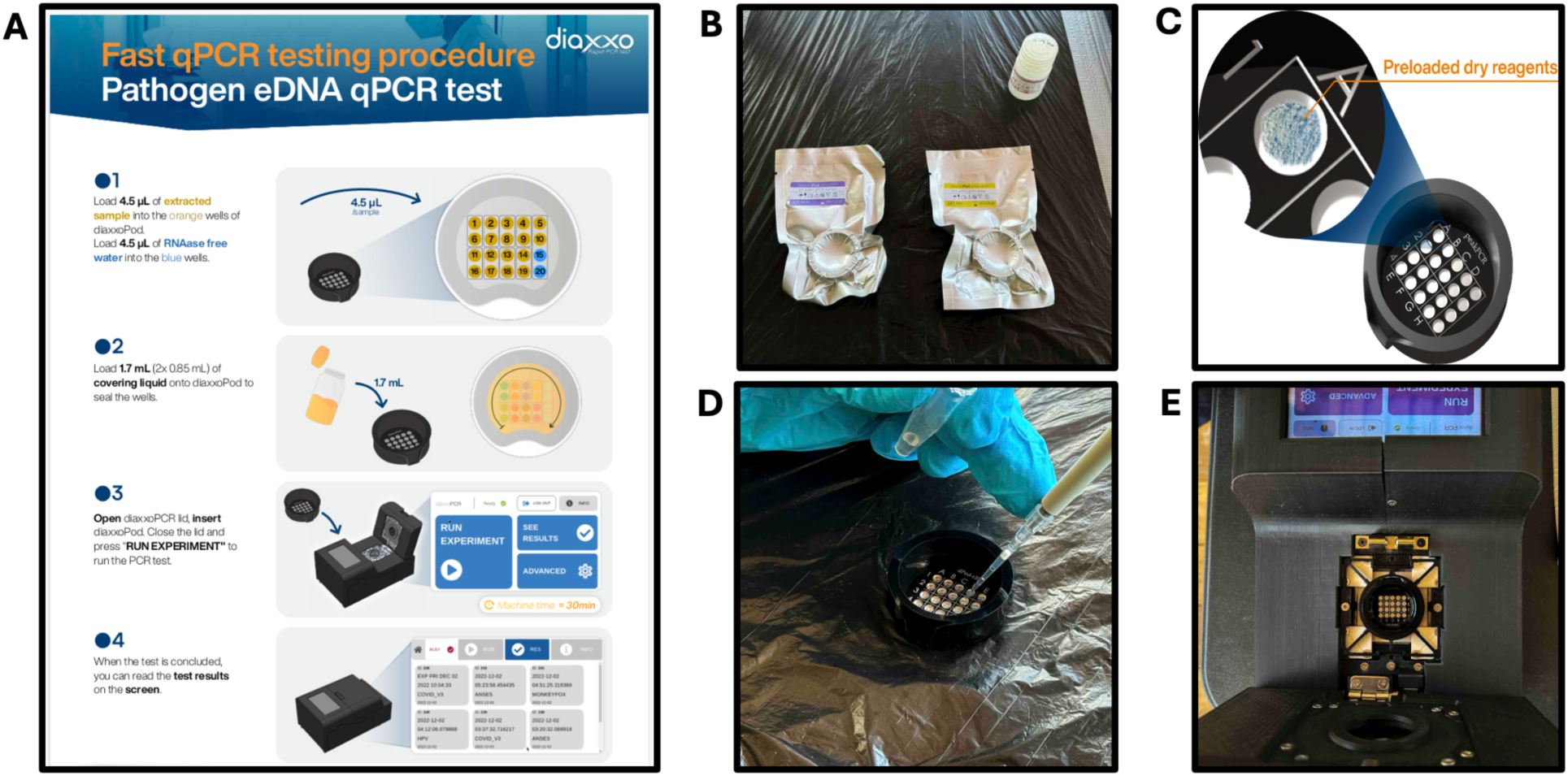
Portable qPCR setup using the Diaxxo system for remote, in situ, parasite detection from environmental DNA (eDNA) samples. (A) Schematic overview of the Diaxxo qPCR procedure, designed for fast and field-deployable detection of pathogens and parasites. The system can process up to 20 reactions simultaneously. (B) Vacuum-packed pods containing lyophilized mastermix and primers in each well, ensuring ease of transport to remote locations without the need for cold chain logistics. The sturdy packaging allows for easy transportation without risk of spills. (C) Close-up schematic of the lyophilized reagents in each well of the PCR pod, pre-loaded and ready for rehydration by adding the eDNA samples. (D) Addition of the eDNA extract into each well, where the sample liquid resuspends the lyophilized reagents and activates the mastermix. A layer of oil is added to prevent cross-contamination between wells, replacing the traditional heated lid used in laboratory thermocyclers. (E) The pod is loaded into the Diaxxo portable qPCR machine for amplification and analysis of the target parasite DNA.

Detection of *Plasmodium* spp. was achieved by targeting the conserved 18S rRNA gene, a region uniformly present across malaria parasites, using a pan-*Plasmodium* primer set adapted from previously validated assays (Bechtold et al., 2023; Stabler et al., 2025). The reaction mixture (proprietary) was prepared one week prior by Diaxxo labs, lyophilized on a qPCR pod (PlasmoPod; Figure 2) and transported via 30 h flight from Switzerland to the test site in Brazil. After which, 4.5 uL of extracted DNA was added directly to each well of the qPCR pod (Figure 2) following the manufacturer’s protocol.

The thermocycling protocol was as follows: an initial incubation at 58 °C for 5 minutes, then an initial denaturation at 91 °C for 3 minutes, followed by an initial extension at 59 °C for 20 seconds. This was succeeded by 45 cycles of 90 °C denaturation for 12 seconds and 59 °C extension for 20 seconds. Images taken at the end of every qPCR cycle were analyzed real-time and cycle threshold (Ct) values were recorded for each sample (Supplementary Material 1), with any value below 35 considered positive for *Plasmodium* spp. Controls included molecular-grade water as negative controls, as well as extraction blanks to ensure no contamination occurred during fieldwork or sample handling. Positive controls were run to confirm the sensitivity and specificity of the assay.

### 2.3 Insect identification from canopy rafts

In parallel with the eDNA sampling, canopy rafts integrating light traps and sticky traps were deployed by drones on the top of forest canopies to survey insect biodiversity (Figure 1 D-F; Geckeler et al., 2025). Four canopy rafts with traps were deployed in the afternoon and left in place for 12 hours before retrieval at dawn the following morning. Upon retrieval, insect specimens were sorted and identified under a microscope, with particular attention to mosquito species, especially those belonging to the *Anopheles* genus, which are known vectors for malaria transmission. Each specimen was carefully removed from the traps using sterile tweezers and placed in Petri dishes for further processing, including imaging and DNA barcoding. Insect specimens were ultimately preserved in ethanol.

DNA extraction was performed via the non-destructive HotSHOT (hot sodium hydroxide and Tris) method (Truett et al. 2000). The HotSHOT procedure yields PCR-ready DNA when using very small quantities of tissue (Gutiérrez-López et al., 2015). Small insects (< 1 cm) were immersed directly into the lysis buffer and tissue from big insects (> 1cm long) was obtained by separating the tibia and tarsomere from the hind legs of each individual using sterile blades. DNA extraction was carried out using 40 μl of lysis buffer (25 mM NaOH, 0.2 mM EDTA, pH 8) and then incubated at 65 °C for 18 minutes followed by 98 °C for 2 minutes. Thereafter, 40 μl of neutralization solution (40 mM Tris-HCl) was added to each sample.

The LCO1490 and HCO2198 (Folmer et al. 1994) primer pair was used to amplify a 658 bp fragment of the COI gene. Each primer contained an additional 13 bp tag on the 5’-end for multiplexing numerous samples for sequencing (Srivathsan et al. 2019). PCRs were performed using Cytiva PuRe Taq Ready-To-Go PCR beads in 0.2 ml tubes (8 tube strips) with 2µl of template DNA, 0.5 µl each of 10 µM tagged primers and 22 µl of molecular grade water to complete a final volume of 25 μl. PCR products were pooled (1 µl per reaction) and purified using AMPureXP beads at 0.75× ratio. The cleaned-up pool was then used for the sequencing library preparation following Srivathsan et al. (2019). The library was sequenced on an Oxford Nanopore MinION device. Consensus DNA barcodes were obtained with ONTbarcoder v2.1.3 (Srivathsan et al., 2024) and subsequently compared to the BOLD database for taxonomic assignment.

### 2.4 Acoustic detection of primates

Passive acoustic monitoring was conducted using AudioMoth recorders (Hill et al. 2019) integrated into the canopy rafts (Figure 1F) to capture vocalizations of animal species living in the forest for biodiversity assessment (Supplementary Figure 1). The recorders were programmed to capture sound continuously using a sampling rate of 46 kHz, and were processed using BirdNET Plus Analyzer (Kahl et al., 2021) with a custom classifier developed by Rainforest Connection that included Howler monkeys in its training. Although seven species of primates are found in the area (IUCN 2024), howler monkeys were the only primates included in the classifier’s database. Given this limitation of the supervised audio species identification, we also performed a manual extraction of calls potentially belonging to primates, which were then validated by a local indigenous scientist from Inhaa-be. Since howler monkeys are known to be potential reservoirs for *Plasmodium* spp., the acoustic data were then cross-referenced with the qPCR results to highlight any correspondence between the presence of howler monkeys and *Plasmodium* spp. DNA in the same area.

## 3. Results

### 3.1 Rapid field detection of *Plasmodium* spp

The entire process, from drone-based eDNA sampling to qPCR results generation, took an average of 1.5 hours per assay (approximately 30 minutes for drone flight, 20 minutes for DNA extraction, and 40 minutes for qPCR). qPCR analysis of eDNA samples from the canopy confirmed the presence of *Plasmodium* spp. in one canopy swab sample. Amplification curves from wells 13 and 18, which were PCR duplicates, produced consistent Ct values of 28.7 and 29.23 (Figure 2), respectively, confirming the positive detection of *Plasmodium* spp. in the sample collected from an area with documented howler monkey activity. The positive control in wells 15 and 20 amplified as expected, with Ct values of 30.82 and 31.11, while the negative control showed no amplification. These results indicate the presence of *Plasmodium* spp. with a low to moderate parasite load in one of the twelve canopy eDNA samples (Figure 3).

**Figure 3.**
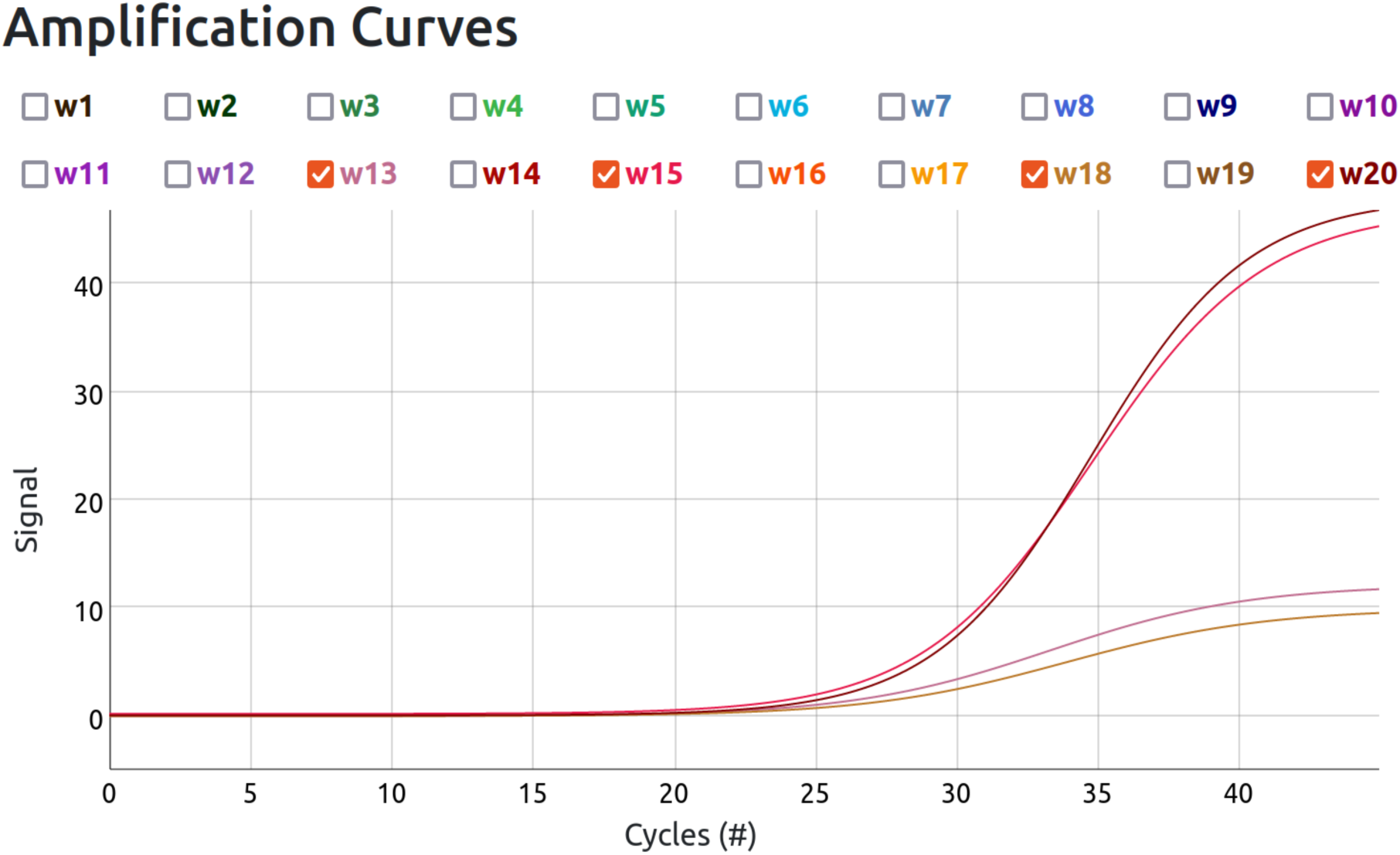
Amplification curves of the surface swab eDNA sample S_005 for the *Plasmodium* spp. assay, showing detection in wells 13 and 18 with Ct values of 28.7 and 29.23, respectively (lower curves). The positive and negative control duplicate assays, loaded with molecular-grade water, are shown in wells 15 and 20 with Ct values of 30.82 and 31.11, respectively (upper curves). These results confirm the detection of *Plasmodium* spp. in the canopy eDNA sample while validating the reliability of the qPCR assay.

### 3.2 Acoustic confirmation of Howler monkey presence

Vocalizations belonging to the Howler monkey (*Alouatta seniculus*) were identified in the recordings of raft number 3 (Figure 1A). Although the quality of segments automatically detected by the classifier was not sufficient for validation, manually extracted audio was confirmed to belong to both adult and infant individuals of *Alouatta seniculus* (Supplementary Audio 1A, B). The overlap between the presence of howler monkeys and positive *Plasmodium* detections suggests that these primates may serve as reservoirs for the parasite. This ecological insight highlights the potential role of primates in maintaining zoonotic disease cycles in tropical ecosystems.

### 3.3 Mosquito surveillance

Visual identification and DNA barcoding of the captured insects revealed a diverse range of species, with *Lepidoptera* (moths) being the most abundant (Supplementary Figure 3). A total of 45 insect species were identified. However, none of the captured insects were known malaria vectors. This result aligns with observations made during the sorting process, where no mosquitoes were found in the sticky traps.

## 4. Discussion

Understanding the intricate ecological relationships in the Amazon rainforest is essential for unraveling the dynamics of zoonoses like malaria (Figure 4). More importantly, real-time detection in the canopy can feed directly into public-health decision-making, providing early warning of parasite presence upstream of human cases. Advanced technologies such as robot-based eDNA sampling, in-situ qPCR testing, and drone-based surveillance have opened new avenues for assessing these complex interactions. By using drones to collect and portable qPCR assays to analyze eDNA from the forest canopy, we successfully detected *Plasmodium* spp. in remote habitats, providing insights into parasite-host interactions that would have otherwise remained undetected. This approach offers a more rapid and comprehensive method for monitoring zoonotic parasites, significantly reducing the need for traditional, labor-intensive sampling and direct wildlife handling.

**Figure 4.**
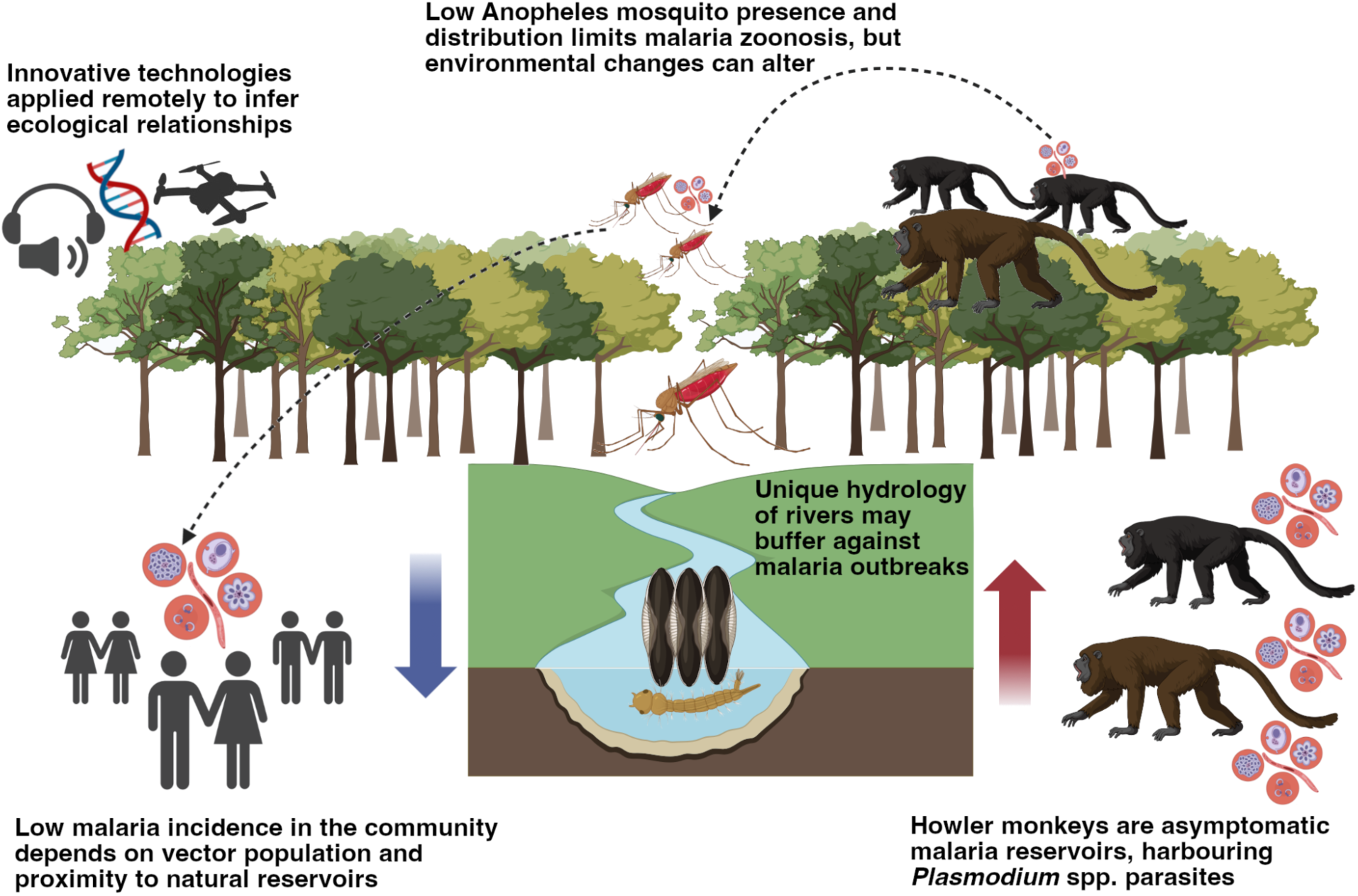
This illustration highlights the ecological relationships inferred using innovative technologies, including drone-based eDNA sampling, portable qPCR assays for *Plasmodium* detection, DNA barcoding of insect traps, and audiometry to confirm howler monkey presence. Our results show no detection of *Anopheles* mosquitoes, consistent with known data that howler monkeys (*Alouatta seniculus*) act as natural reservoirs for *Plasmodium* spp. in the Amazon. We propose that the current low malaria risk in the region is due to the scarcity of *Anopheles* mosquitoes, underscoring the importance of enhanced biodiversity monitoring for proactive disease prevention. Images adapted from Biorender.

The detection of *Plasmodium* eDNA in canopy swabs suggests parasite presence in arboreal wildlife, likely through bodily fluids or fecal matter. This finding aligns with prior studies identifying howler monkeys (*Alouatta seniculus*) as asymptomatic carriers of *Plasmodium* spp. in Latin America (Abreu et al., 2019). However, the absence of *Anopheles* in our light-based canopy traps, despite likely underestimating true vector abundance, somewhat aligns with the low incidence of malaria in nearby human populations. Canopy entomological surveillance more commonly employs CO₂-baited traps or human landing catches (which are generally more effective for host-seeking *Anopheles* spp.), so our light trap may have underrepresented true vector abundance (Le Breton et al., 2025). As such, while this pattern is intriguing, these findings only suggest low vector densities rather than confirm them.

The hydrological characteristics of blackwater rivers in the sampling area, like the Rio Negro, create suboptimal breeding conditions for *Anopheles* species, including *Anopheles darlingi*, the primary malaria vector in the region (Fonseca et al., 2022; 2023). These environmental factors, coupled with low vector densities, may serve as natural barriers preventing zoonotic spillover of malaria parasites into nearby human populations. This also suggests that malaria transmission in this region may not follow the typical vector-host-parasite model, where vector suppression, rather than parasite absence, can drive malaria transmission dynamics, and spillover could occur only under specific environmental conditions. Additionally, shifting transmission risk in response to climate change, with rising temperatures and altered rainfall patterns, may allow the malaria-carrying mosquitoes to thrive in once unsuitable areas. Ongoing environmental changes, such as deforestation and global warming, could further exacerbate this, potentially increasing mosquito populations and leading to future malaria outbreaks (Mereta et al., 2013; Estifanos et al., 2025).

The integration of eDNA sampling with drone technology and portable qPCR assays provides a powerful framework for monitoring species interactions and parasite dynamics in previously inaccessible areas of the rainforest. The absence of mosquitoes in canopy traps, coupled with the presence of *Plasmodium* in eDNA swabs, underscores the importance of vector distribution in shaping disease dynamics. This approach complements traditional surveillance methods and offers early warnings of disease presence, enabling the monitoring of multiple parasites with fewer resources. The rapid nature of our workflow with drone-based eDNA sampling, combined with portable qPCR assays, highlights the transformative potential of portable, real-time and non-invasive eDNA technology for parasite surveillance, especially in remote and resource-limited settings.

In 2023 alone, Brazil lost approximately 3.07 million hectares of natural forest (Global Forest Watch, 2024). Such accelerating deforestation rates in the Amazon heighten the risk of zoonotic spillover by fragmenting wildlife habitats and altering vector-host dynamics. Effective disease prevention must therefore consider the interconnectedness of local communities, wildlife, and their ecosystems. Our findings highlight the importance of biodiversity conservation as a key pillar of disease mitigation, in alignment with the One Health framework (Arisco et al., 2024; Erkyihun et al., 2022). Raising awareness about biodiversity and zoonotic disease risks is vital to safeguarding human and wildlife health (Kilpatrick et al., 2017). Future research should focus on mapping vector species distributions, understanding seasonal variations in vector-host interactions, and monitoring primate populations. These efforts are critical for predicting how environmental changes will influence malaria transmission in the Amazon and other biodiverse regions. Adopting innovative sampling methods and fostering cross-sector collaboration will lead to more effective strategies for protecting human and animal populations from emerging infectious diseases.

In conclusion, our findings suggest that howler monkeys (*Alouatta seniculus*) and potentially other non-human primates may serve as asymptomatic reservoirs for *Plasmodium* spp. in parts of the Amazon. However, the limited presence of *Anopheles* mosquitoes remains the primary ecological barrier preventing malaria transmission to humans in these regions.

Maintaining forest integrity and minimizing human disturbances are, therefore, crucial for mitigating zoonotic malaria risks. By establishing a rapid, drone-assisted parasite detection workflow with in-situ qPCR readout, we demonstrate a field-deployable platform capable of triggering proactive vector-control and outbreak-response measures—fulfilling a critical need in remote, high-risk regions. As environmental changes such as deforestation and climate shifts accelerate, the risk of malaria spillover could increase. Addressing these risks will require proactive efforts to preserve intact ecosystems, monitor vector populations, implement early detection systems, and promote biodiversity conservation as a core strategy for reducing zoonotic disease emergence.

## 5. Author Contributions

The study was supervised by S.M. and K.D. S.K, C.G. and G.S. deployed the drones for eDNA collection. A.D., K.D., M.J., M.L., Y.C.A.I. and L.M. conducted the eDNA experiments in the field; Y.C.A.I., and L.M. performed the data analyses. L.M., M.G., and T.S. built the portable DiaxxoPCR machine and designed the qPCR assay. F.B.T. and C.M. designed and deployed the canopy traps. P.P.S., N.D.F.S. and M.A.G.H. processed the specimens collected from the canopy rafts and analysed the data. F.S. performed the audio analysis and analysed the results, with validations by M. M. Manuscript draft and figures were prepared by Y.C.A.I., with inputs from J.J.M.C, L.M., S.M., and K.D. All authors contributed to the manuscript edits, and approved the final version for publication.

## 6. Declarations

This study was funded by the Rütli-Stiftung, the ETH Foundation, the XPRIZE Foundation, and the Alana Foundation via the participation of the authors in the XPRIZE Rainforest competition. S.M., S.K., and C.G. were supported by the Swiss National Science Foundation through the Eccellenza grant (grant number 186865). All data are available in the main text and supplementary data. Competing interests: M.G., L.M., G.M. and T.S. are employed by, and shareholders of, the ETH Zurich spin-off company—Diaxxo AG—the organization that produces the DiaxxoPCR device. The Method and UAV for collecting environmental DNA is protected for commercial use under the patent WO2024246233A1. K.D. is the CEO of SimplexDNA. All other authors declare that they have no competing interests.

## 9. Supplementary

Supplementary Audio 1A and B. *Alouatta seniculus* adult and infant field audiomoth recording.

**Supplementary Figure 1.**
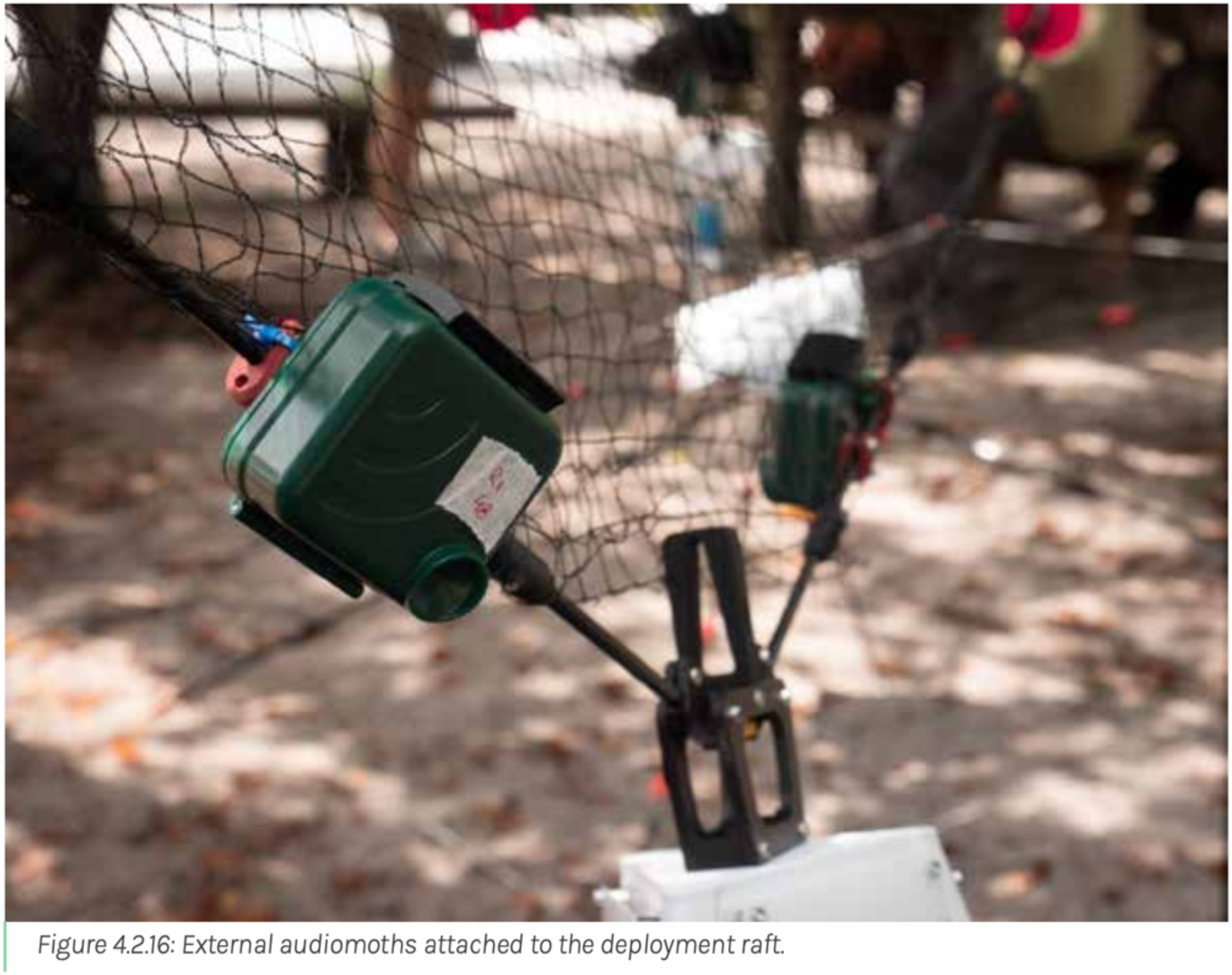
External audiomoths attached to the deployment raft.

**Supplementary Figure 2.**
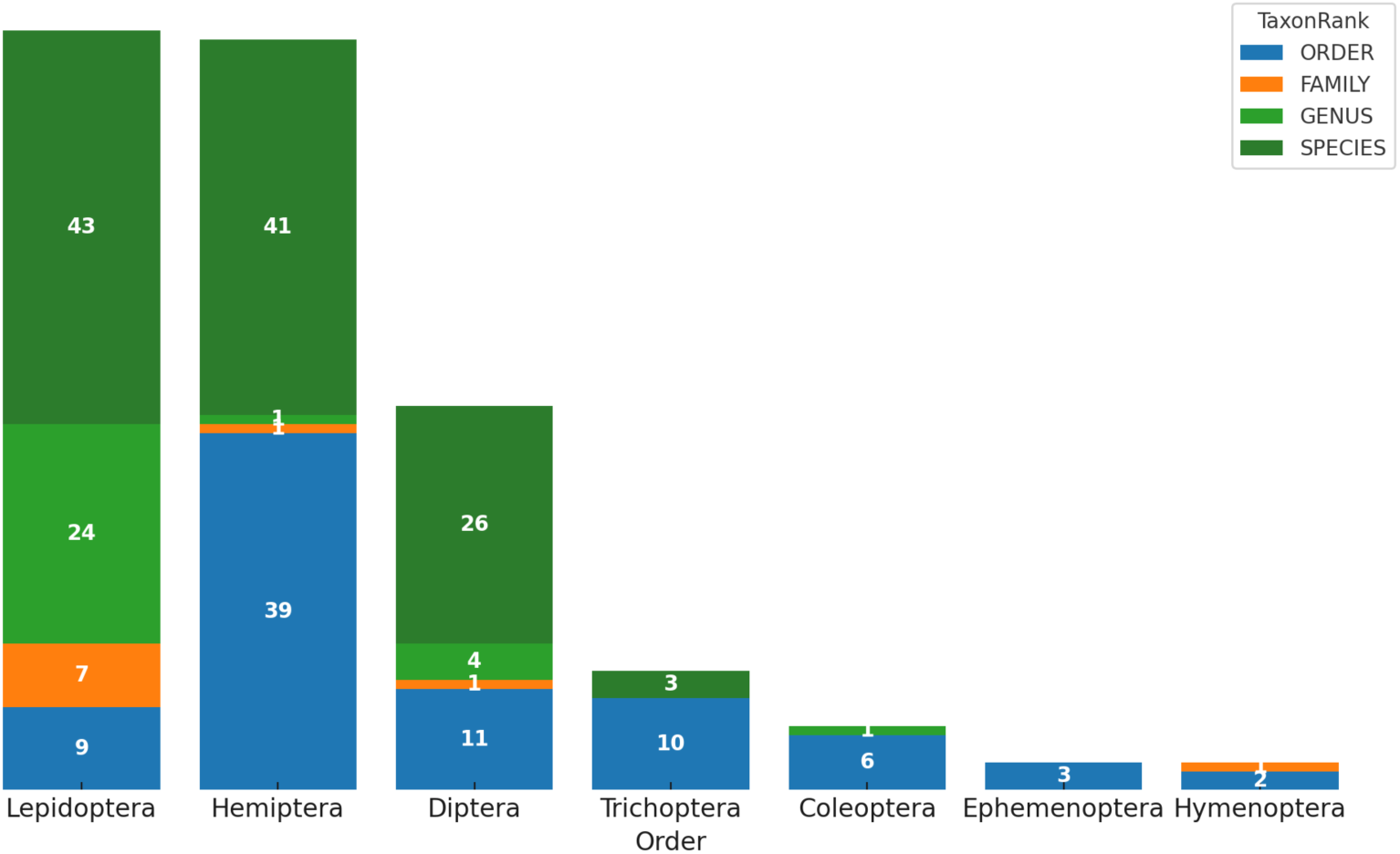
DNA barcoding of captured insects revealed a diverse range of species, with Lepidoptera (moths) being the most abundant (n=83). A total of 45 species were identified, though no malaria vectors were found, consistent with the absence of mosquitoes in the sticky traps.

